# RAS and PP2A activities converge on epigenetic gene regulation

**DOI:** 10.1101/2022.05.11.491459

**Authors:** Anna Aakula, Mukund Sharma, Francesco Tabaro, Henrik Honkanen, Matthieu Schapira, Cheryl Arrowsmith, Matti Nykter, Jukka Westermarck

## Abstract

RAS-mediated human cell transformation requires inhibition of the tumor suppressor Protein Phosphatase 2A (PP2A). Both RAS and PP2A mediate their effects by phosphoregulation, but phosphoprotein targets and cellular processes in which RAS and PP2A activities converge in human cancers have not been systematically analyzed. Here, based on mass spectrometry phosphoproteome data we discover that phosphosites co-regulated by RAS and PP2A are enriched on proteins involved in epigenetic gene regulation. As examples, RAS and PP2A co-regulate the same phosphorylation sites on HDAC1/2, KDM1A, MTA1/2, RNF168 and TP53BP1. Mechanistically, we validate co-regulation of NuRD chromatin repressor complex by RAS and PP2A. Consistent with their known synergistic effects in cancer, RAS activation and PP2A inhibition resulted in epigenetic reporter de-repression and activation of oncogenic transcription. Notably, transcriptional de-repression by PP2A inhibition was associated with increased euchromatin and decrease in global DNA methylation. Further, targeting of RAS- and PP2A-regulated epigenetic proteins decreased viability of KRAS-mutant human lung cancer cells. Collectively the results indicate that epigenetic protein complexes involved in oncogenic gene expression constitute a significant point of convergence for RAS hyperactivity and PP2A inhibition in cancer. Further, the results provide a rich source for future understanding of phosphorylation as a previously unappreciated layer of regulation of epigenetic gene regulation in cancer, and in other RAS/PP2A-regulated cellular processes.

## INTRODUCTION

*RAS* genes (*HRAS, KRAS*, and *NRAS*) comprise the most frequently mutated oncogene family in human cancer, accounting for 3.5 million new cases yearly, worldwide (Prior, Hood et al., 2020). RAS-mediated human cell transformation is preceded by cell immortalization which is caused by loss of tumor suppressors such as TP53, RB1 or CDKN2 (Minna, Roth et al., 2002). Hyperactivation of RAS signaling is however not alone sufficient for malignant transformation of immortalized human cells, but requires simultaneous inhibition of the phosphatase activity of the tumor suppressor Protein Phosphatase 2A (PP2A) (Hahn, Dessain et al., 2002, Rangarajan, Hong et al., 2004, Sablina, Hector et al., 2010, Sato, Larsen et al., 2013, Tian, Doerig et al., 2018, Yu, Boyapati et al., 2001). PP2A inhibition and RAS mutations also significantly synergize in predicting poor overall survival of cancer patients across the TCGA pan-cancer data (Kauko, Laajala et al., 2015). Moreover, reactivation of the tumor suppressor activity of PP2A efficiently inhibits RAS-driven tumorigenesis (Liu, Gu et al., 2015, Saddoughi, Gencer et al., 2013, Sangodkar, Perl et al., 2017), and synergize with pharmaceutical targeting of RAS downstream effector MEK (Kauko, O’Connor et al., 2018). Thus, understanding the mechanistic basis of the synergism between PP2A inhibition and RAS activity could provide significant novel opportunities for targeting RAS-dependent cancers.

PP2A comprises a family of trimeric protein complexes that counter-balance kinase-mediated phosphorylation throughout cell signaling networks (Fowle, Zhao et al., 2019). The PP2A trimers are composed of a scaffolding PP2A-A subunit, a catalytic C subunit, and one of the alternative substrate determining B-subunits. In about 10% of human cancers PP2A is inhibited by genomic mutations, but the most prevalent mechanism for PP2A inhibition in cancer is overexpression of one of the numerous oncogenic PP2A inhibitor proteins such as CIP2A, PME1 or SET (Kauko & Westermarck, 2018) (Fig. 1A). Several downstream effectors/kinases of RAS are identified as PP2A targets, but it is not yet fully elucidated what the cancer-relevant cellular processes in which RAS and PP2A activities converge. Indeed, PP2A has been shown to regulate for example RAF-MEK-ERK and PI3K-AKT pathways (Fowle et al., 2019, Kauko & Westermarck, 2018, Sablina et al., 2010), but whether there are processes beyond kinase signaling that are relevant to RAS/PP2A co-operation in cancer is poorly understood. Interestingly, a recent phosphoproteome analysis revealed that PP2A inhibition and RAS activity regulate highly overlapping phosphoproteins (Kauko et al., 2015). However, the functional relevance of these findings has not been studied yet.

**Figure 1.**
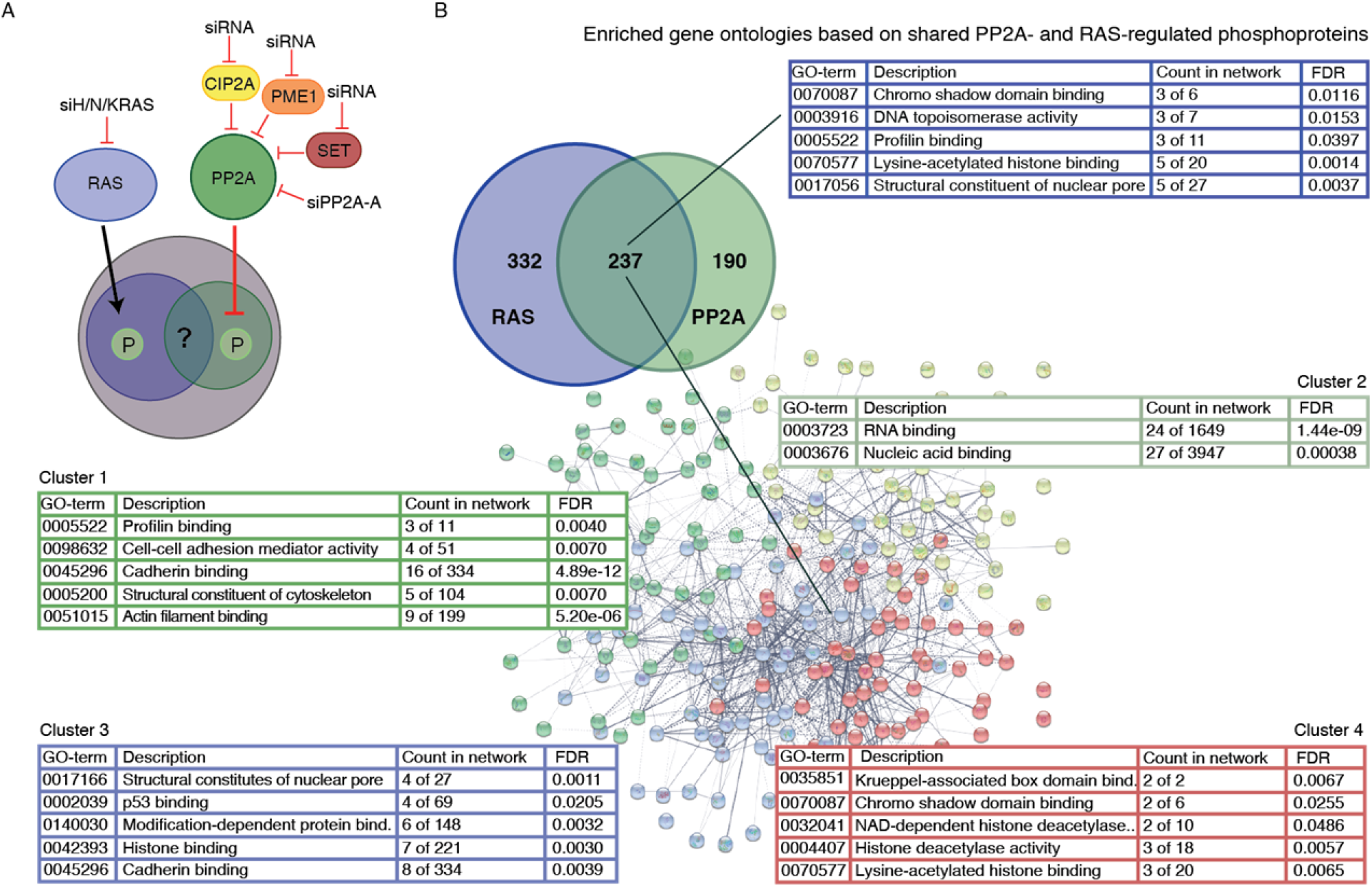
Enriched gene ontologies based on shared RAS- and PP2A-regulated phosphoproteins. **A**. The schematic presents the conducted phosphoproteomics set-up, utilizing siRNAs for RAS (H/K/N), PP2A-A and PP2A inhibitory proteins (CIP2A, PME1 and SET). **B**. The figure shows the identified overlap between RAS- and PP2A-regulated phosphoprotein target, as well as the enriched gene ontologies (GO terms) related to these.

Gene expression in multicellular organisms is regulated through various epigenetic processes involving for example insertion or removal of chemical tags on the nucleotides and histones (Miller & Grant, 2013). Importantly, both epigenetic gene silencing of tumor suppressors, as well as increased transcription of oncogenic genes contribute to cancer initiation, progression, and therapy resistance (Baylin & Jones, 2016, Quagliano, Gopalakrishnapillai et al., 2020). DNA methylation is the best characterized epigenetic mechanism mediating gene repression. On the other hand, mechanisms that impact nucleosomes via covalent histone modifications leading to open chromatin state (euchromatin) are well established for oncogenic transcription. Strategies to impact epigenetic gene regulation might therefore be useful in cancer prevention and therapy (Cheng, He et al., 2019, Quagliano et al., 2020).

While the mechanism of action of epigenetic proteins is extensively studied, and loss-of-function approaches demonstrated that many have a role in cancer (Baylin & Jones, 2016, Laugesen & Helin, 2014), generally very little is known about how phosphorylation-dependent oncogenic signaling regulates their activities (Trevino, Wang et al., 2015). One of the major epigenetic complexes involved in cancer is the Nucleosome Remodeling and Deacetylase complex (NuRD) which functions by two different enzymatic activities; the ATP dependent nucleosome remodeling through CHD3/4/5 and deacetylation of histone tails through HDAC1/2 (Laugesen & Helin, 2014). HDAC1/2 is one of the rare epigenetic proteins known to be subject to phosphoregulation by kinase/PP2A balance (Bahl & Seto, 2021). However, beyond HDAC1/2, (de)phosphoregulation of the other members of the NuRD complex is poorly understood. DNA methyltransferases are another group of cancer relevant epigenetic writer proteins. DNMT1 is known to be regulated by phosphorylation at S143 by AKT, resulting in increased protein stability (Esteve, Chang et al., 2011). However, there is no published information about the potential role of phosphatases regulating DNMT1. Further, although the potential for methyltransferase DOT1L as a therapy target in KRAS mutant cancers was recently demonstrated (Liu, Liu et al., 2021), currently there is neither information whether RAS regulates DOT1L phosphorylation, nor any indications for the role of phosphatases on DOT1L phosphoregulation.

Here, we have addressed the open question of convergence of RAS- and PP2A-mediated phosphoregulation in cancer by using previously published phosphoproteome datasets in which RAS proteins and PP2A complexes were targeted by siRNAs (Kauko, Imanishi et al., 2020, Kauko et al., 2015) (Fig. 1A). The results demonstrate that epigenetic gene regulation is particularly enriched among the cellular processes co-regulated by RAS- and PP2A-mediated phosphorylation. This is due to numerous, but previously unidentified, RAS- and PP2A-regulated phosphosites in epigenetic proteins implicated in cancer. Functionally we validate the role of both RAS activity and PP2A inhibition in oncogenic transcription, and demonstrate the first evidence for PP2A in regulation of DNA methylation and chromatin remodeling. Together with the demonstration of the oncogenic role of several RAS/PP2A-regulated epigenetic targets, as well as the role of PP2A inhibition in HDAC inhibitor resistance, these data unveil an unprecedented role for PP2A/RAS-mediated (de)phosphorylation in epigenetic complexes involved in oncogenic transcription.

## RESULTS

### Systematic analysis of phosphosites co-regulated by RAS- and PP2A

To comprehensively map the phosphoproteins co-regulated by PP2A and RAS, we combined data from two recent phosphoproteome studies, in which either all three forms of RAS (HRAS, KRAS and NRAS)(Kauko et al., 2015), or the PP2A scaffold protein PP2A-A, or the PP2A inhibitor proteins CIP2A, PME1, and SET (Kauko et al., 2020), were depleted by siRNAs (Fig. 1A, Table S1). For RAS and the PP2A inhibitory proteins we included the phosphosites that were dephosphorylated in the siRNA transfected cells, whereas for PP2A-A we included the phosphosites that had increased phosphorylation (see Materials and Methods for data filtering). The resulting RAS/PP2A phosphoproteome consisted of 1,518 unique phosphosites in 749 proteins. RAS inhibition resulted in dephosphorylation of 725 phosphosites in 427 proteins, whereas the corresponding values for PP2A-A and the PP2A inhibitor proteins were 274/195 and 875/441, respectively (Table S2). As a clear indication for convergence of RAS and PP2A activities on phosphoproteome regulation, altogether 270 distinct phosphorylation sites on 237 proteins were found to be co-regulated both by RAS and PP2A targeting (Fig. 1B). Interestingly, when assessing the overlap of the regulated phosphosites between RAS and PP2A modulations, sites dephosphorylated by RAS inhibition overlapped more frequently with sites dephosphorylated by PP2A inhibitor protein inhibition, than with sites regulated by PP2A inhibition (Fig. S1). This can be explained by the notion that both RAS inhibition and PP2A reactivation inhibit phosphorylation of sites that are constitutively phosphorylated in cancer cells, and based on the recent model that the majority of cellular phosphosites are exclusively dominated by either phosphatase activation or inhibition (Kauko et al., 2020). The overlap between RAS and PP2A inhibitor protein SET was particularly notable (Fig. S1A,B). This could explain very potent antitumor effects of SET inhibition in RAS-driven tumorigenesis (Liu et al., 2015, Saddoughi et al., 2013). In addition to 237 proteins in which at least one phosphorylation site was co-regulated by both RAS and PP2A (Fig. 1B), RAS and PP2A co-regulated phosphorylation of 57 overlapping proteins but in these proteins the RAS and PP2A regulated sites were not identical (Table S2). Collectively these analyses demonstrate a clear convergence of RAS and PP2A-mediated phosphoregulation, both at the level of individual phosphorylation sites, but also at the level of proteins.

To identify cellular processes that would be governed by the convergence of RAS- and PP2A-mediated phosphoregulation, we analyzed enriched gene ontologies (GO) based on the 237 proteins in which there was at least one phosphosite regulated by both RAS and PP2A (Fig. 1B). The STRING database (Szklarczyk, Gable et al., 2021) analysis revealed a clear enrichment of GOs related to epigenetic and transcriptional gene regulation (Fig. 1B, GO terms 0070087, 0003916, and 0070577). To increase the resolution of the analysis, we performed an analysis in which the shared RAS/PP2A phosphotargets were divided into four clusters (Fig. 1B). Whereas cluster 1 was mostly associated with cytoskeleton and cell adhesion, and cluster 2 with nucleic acid binding, clusters 3, and especially 4, revealed a very strong association of the target proteins with histone modifications and chromatin remodeling (Fig. 1B). While recent data validate the critical role for PP2A in regulating transcriptional elongation (Huang, Jee et al., 2020, Vervoort, Welsh et al., 2021), and epigenetic gene regulation has important role in RAS-mediated oncogenesis (Vaz, Hwang et al., 2017), the role of PP2A and RAS in phosphorylation-dependent regulation of epigenetic complexes is very poorly -understood. Based on these notions, we focused our downstream analysis on RAS and PP2A convergence on epigenetic gene regulation and transcription.

### PP2A- and RAS-mediated phosphorylation converge on epigenetic complexes

Interestingly, many epigenetic RAS/PP2A phosphotargets were found to constitute protein complexes with each other (Fig. 2A). The most apparent examples were the NuRD, DNMT1 and DOT1L complexes (Fig. 2A). We postulate that RAS/PP2A signaling can potentially regulate DNA methylation, histone methylation, and histone deacetylation via phosphorylation of these complexes (Trevino et al., 2015) (Fig. 2A). Consistent with the convergence model, most protein members of these epigenetic complexes were regulated by both RAS and PP2A (Fig. 2A), both at the level of individual phosphosites, but also at the level of proteins (Fig. 2B, S2). Naturally some epigenetic factors were also found to be regulated by either RAS or PP2A only (Fig. 2B, S2). In the case of PP2A-regulated targets, the evidence for direct PP2A-mediated dephosphorylation was strengthened by the identification of putative binding motifs for PP2A B-subunit B56 in many of these proteins (Table S3). Based on previous evidence, that B56 binding motif containing proteins can act as scaffolds for the recruitment of the other complex proteins for PP2A-mediated dephosphorylation (Hertz, Kruse et al., 2016), the majority of the individual PP2A target proteins from our data could become accessible for PP2A-mediated phosphoregulation.

Structural evaluation of the phosphorylation sites on selected RAS/PP2A targets provided clues about their potential functional importance. Using the RNF168 paralog, RNF169-Histone 2 complex, as the model structure (PDB), the RAS/PP2A-regulated RNF168 serine 481 was found adjacent to arginine 466 that is critical for histone binding (Fig. 2C). On the other hand, PP2A-regulated serine 714 on DNMT1 is located at the base of the loop that must relocate for DNA binding (Fig. 2C).

We further characterized the oncogenic potential of selected PP2A/RAS regulated epigenetic target proteins. To this end, we targeted the selected proteins with three siRNAs per gene, in three KRAS mutant non small cell lung cancer (NSCLC) cell lines, A549, H358 and H460 (Fig. S3A). In parallel, H358 cells were treated with chemical inhibitors for DNMT1, bromodomain proteins, and HDAC1/2 (Fig. S3B). Collectively the results show a significant role for most of the epigenetic RAS/PP2A target proteins for NSCLC cell viability.

In summary, these data demonstrate convergence of PP2A- and RAS-mediated phosphoregulation on cancer relevant epigenetic protein complexes.

### PP2A and RAS regulate the NuRD complex

Based on the phosphoproteome data (Fig. 2), we proceeded to validate the impact of PP2A and RAS on selected epigenetic factors, and especially on chromatin recruitment of the NuRD complex as it showed very high level of PP2A/RAS-mediated phosphorylation regulation (Fig. 2A).

**Figure 2.**
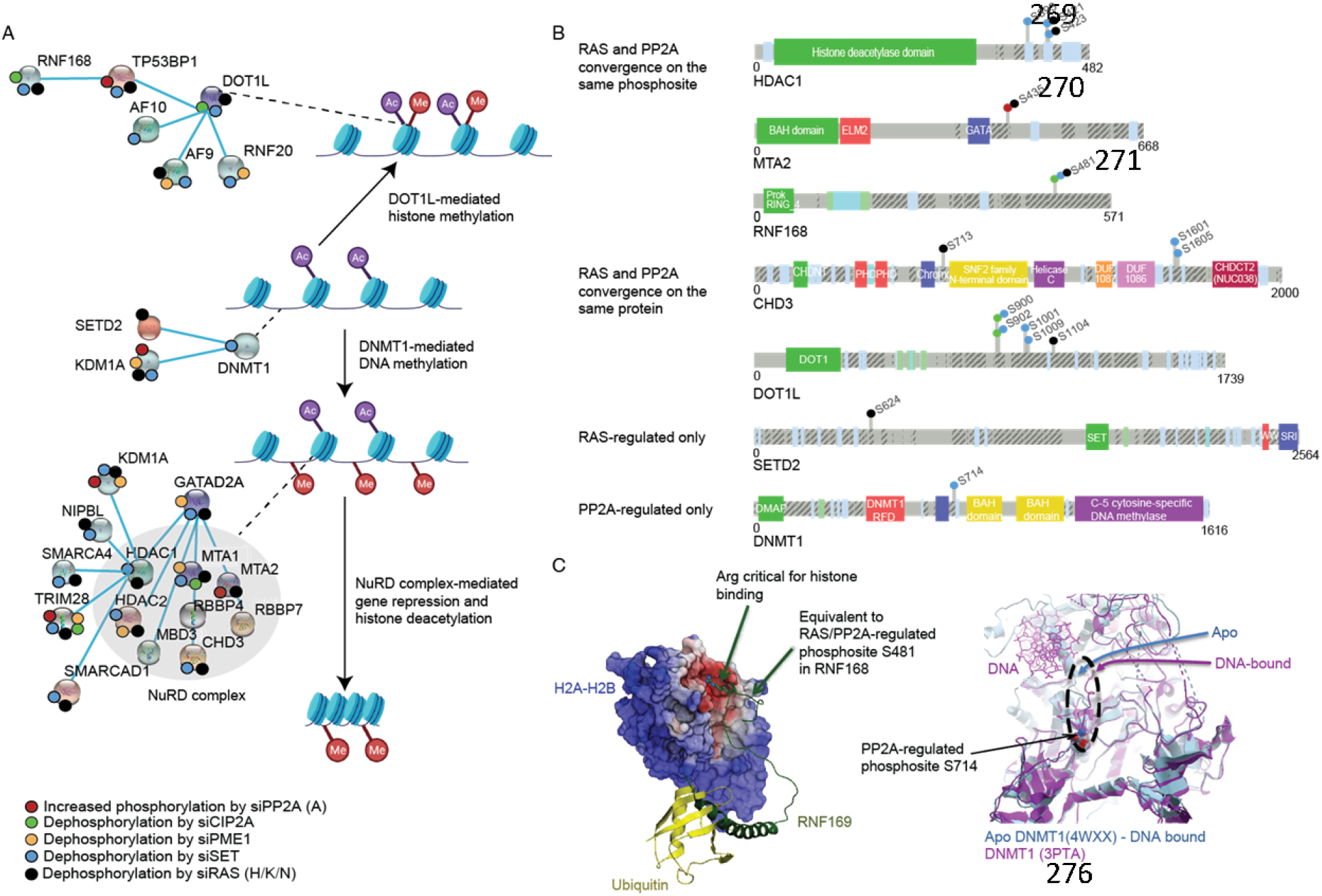
PP2A- and RAS-mediated phosphorylation converge on epigenetic complexes. **A**. The schematic presents some of the PP2A and RAS-regulated proteins involved in opening and closing of the chromatin. The small colored dots on each of the proteins indicate the identified change in phosphorylation in mass spectrometry upon siRNA treatments of either PP2A-A, its inhibitory proteins (CIP2A, PME1 or SET) or by RAS (H/K/N) **B**. The figure highlights selected PP2A-and RAS-regulated proteins, and their respective (de)phosphorylation sites regulated by these. The small colored dots again indicate the identified change in phosphorylation, upon siRNA treatments. **C**. Structural modeling of RAS/PP2A-regulated phosphosites on selected epigenetic proteins.

CHD3 is a member of the NuRD complex, and it supports viability of KRAS-mutant lung cancer cell lines (Fig. S3). Based on Phosphosite Plus database (www.phosphosite.org) CHD3 is phosphorylated on 54 distinct serines or threonines but there are no reports about the functional relevance of any of these phosphosites. In our data, RAS inhibition resulted in dephosphorylation of S713 (Fig. 3A and S2), whereas PP2A reactivation by SET inhibition caused dephosphorylation of S1601 and S1605 (Fig. S2, Table S1). Using yeast CHD1 crystal structure (PDB 3MWY) as a model, the RAS target site S713 was located on the unstructured region in the vicinity of the nucleotide binding mediating region between amino acids 761-768 in human CHD3 protein (Fig. 3A). This structural organization was supported by AlphaFold analysis of human CHD3 (Fig. S4A). S713 phosphorylation in KRAS mutant A549 lung cancer cells seems to be critical for the stability of CHD3 protein, as S713A mutation dramatically inhibited CHD3 protein expression, whereas the phosphorylation mimicking mutation S173D resulted in increased protein expression (Fig. 3B). Next, we evaluated the potential impact of PP2A/RAS activities on NuRD complex chromatin recruitment by following intranuclear distribution of HDAC1/2 as central components of the NuRD complex. Notably, PP2A activation, either by siPME1 or by the pharmacological activator DBK1154 (Vervoort et al., 2021), enhanced both nucleoplasmic localization and the chromatin recruitment of HDAC2 (Fig. 3C, D, E and F). Furthermore, the interaction between the PP2A B-subunit B56α and HDAC1 was confirmed by pulldown experiments (Fig. S5A). Consistent with the prominent impact of RAS on NuRD complex phosphorylation (Fig. 2A), RAS inhibition also resulted in increased nucleoplasmic and chromatin retention of HDAC1/2 (Fig. 3G, H). To link the HDAC regulation by PP2A inhibition and RAS activity to their co-operative roles in human cell transformation, we evaluated chromatin recruitment of HDAC1 from human bronchial epithelial cells (HBEC) transformed by serial introduction of short hairpin p53 (p53-), RAS G12V overexpression (KRAS+), as well as overexpression of the viral PP2A inhibitor protein small-t (ST). Consistent with the published results (Sato et al., 2013), only the cells with both RAS activation and PP2A inhibition were able to grow on soft agar as a measure of cellular transformation (Fig. S5B, C). Interestingly, p53 inhibition in HBECs resulted in robust chromatin recruitment of HDAC1 (Fig. 3I, J). However, the chromatin recruitment was reversed by RAS activation and this RAS-elicited response was further stabilized by ST-elicited PP2A inhibition (Fig. 3J). We further asked whether co-operative inhibition of HDAC1/2 chromatin recruitment by RAS activity and PP2A inhibition impacts cellular sensitivity to the pharmacological HDAC inhibition. To this end, PP2A-A was depleted from KRAS mutant H460 cells, and the cells were treated with increasing concentrations of the clinical stage HDAC inhibitor Panobinostat (Fig. S5d). Alternatively, we tested the potential synergy between Panobinostat and PP2A activator DBK1154 (Fig. S5E, F). In both settings we observed that the impact of PP2A on nucleoplasmic/chromatin distribution of HDAC1/2 correlated with the sensitivity of cells to Panobinostat. Decreased chromatin recruitment by PP2A inhibition (Fig. 3H, I) correlated with Panobinostat resistance, whereas increased chromatin recruitment by PP2A activation (Fig. 3C, D) correlated with increased Panobinostat sensitivity.

These results clearly indicate that PP2A/RAS-mediated phosphoregulation also functionally impacts epigenetic complexes in cancer cells. However, understanding of the functional role of each identified phosphorylation site reported here will require extensive validation experiments outside the scope of this resource article.

**Figure 3.**
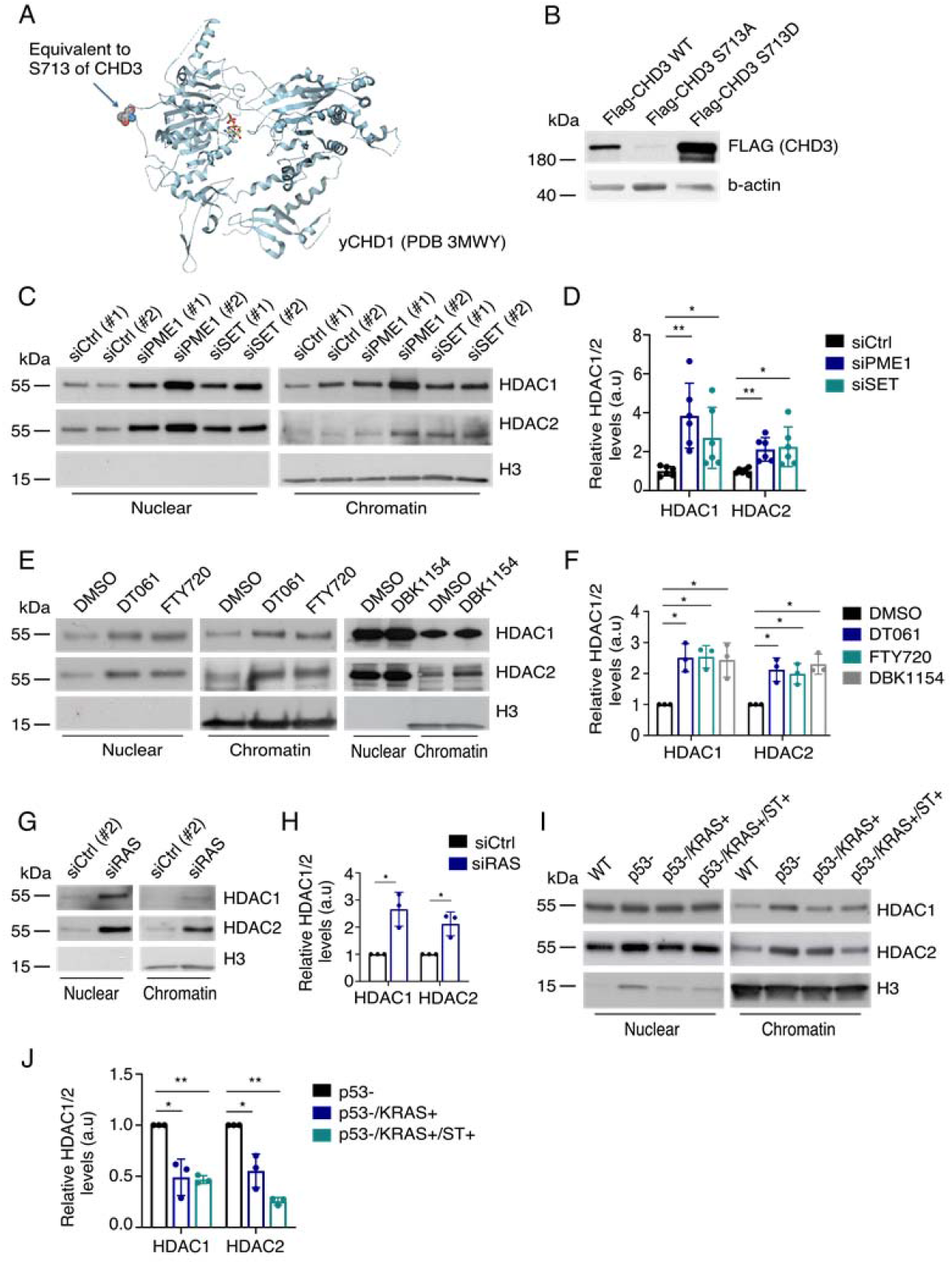
Effects on NuRD complex proteins upon PP2A and RAS modulations. **A**. CHD3 protein and the location of the identified RAS-regulated serine 713. **B**. Western blot analysis of the effect of the S713A and S713D mutations on protein stability upon CHD3 overexpression. **C**. Western blot analysis of chromatin recruitment of HDAC1 and 2 upon siRNA-mediated inhibition of indicated PP2A inhibitory proteins. **D**. Quantification of siPME1 and siSET effect on HDAC1/2 recruitment to chromatin. **E**. Western blot showing the impact of PP2A activation (DT061 and DBK1154) and SET inhibition (FTY720) on HDAC1/2 recruitment to chromatin. **F**. Quantification of DT061, FTY720 and DBK1154 effect, as compared to DMSO, on HDAC1/2 recruitment to chromatin. **G**. Western blot analysis of the impact of RAS inhibition on HDAC1/2 chromatin recruitment. **H**. The quantification of chromatin recruitment upon siRAS from WB in E. **I** and **J**. Chromatin recruitment of HDAC1/2 in the step-wise transformed HBEC cells, as Western blot and quantification, respectively.

### PP2A inhibition and RAS activity de-repress transcription

To address the functional impact of RAS- and PP2A-mediated co-regulation of epigenetic complexes on gene regulation, we employed a previously established stable cell line carrying an epigenetically silenced SFRP1 promoter-GFP reporter (Cui et al., 2014) (Fig. 4A). De-repression of the promoter results in GFP expression that can be monitored by Western blotting or by live cell IncuCyte analysis (Cui et al., 2014). As a technical control, we verified that treatment of cells with the DNMT1 inhibitor 5-aza-2’-deoxycytidine (Decitabine) resulted in increased SFRP1-GFP reporter activity (Fig. 4B). Notably, the reporter was also responsive to pharmacological inhibition of other epigenetic PP2A/RAS phosphotargets, such as HDACs and KDM1A (Fig. S5G, H), further validating its suitability as a surrogate reporter for PP2A/RAS-mediated regulation of epigenetic gene regulation.

To test the impact of RAS/PP2A activities on SFRP1 promoter activity, we used the same siRNA treatments as were used for generating the phosphoproteome data (Kauko et al., 2020, Kauko et al., 2015). Notably, inhibition of PP2A either by siPP2A-A, or by chemical serine/threonine phosphatase inhibitor Okadaic acid, resulted in increased promoter activity (Fig. 4B, C). Related to the endogenous PP2A inhibitory mechanisms, PME1 depletion resulted in further repression of SFRP1-GFP reporter activity, while SET depletion did not have an effect (Fig. 4B, C). The role for PME1-mediated PP2A inhibition in promoting oncogenic transcription was further supported by the increased reporter activity upon transient PME-1 overexpression (Fig. 4D, E). On the other hand, RAS inhibition resulted in significant decrease in SFRP1-GFP reporter activity (Fig. F and G). Downstream of RAS, the effect on reporter activity appears to be at least partly mediated by MEK-ERK MAPK pathway, as MEK inhibitor AZD6244 treatment also resulted in significant reporter activity inhibition (Fig. 4E).

These data indicate that like their opposing roles in oncogenesis (Rangarajan et al., 2004, Sangodkar et al., 2017, Sato et al., 2013, Zhou, Updegraff et al., 2017), PP2A and RAS have opposite roles in epigenetically regulated gene expression.

### Transcriptional profiling of PP2A- and RAS-inhibited cells

To further evaluate the directionality of PP2A -and RAS-mediated gene regulation, we analyzed global gene expression patterns upon PP2A or RAS inhibition by RNA sequencing (RNA-seq). Consistent with the results obtained by the SFRP1-GFP reporter system, PP2A inhibition predominantly resulted in global gene activation, while RAS inhibition predominantly led to gene repression (Fig. 5A and B). Notably, gene set enrichment analysis (GSEA) of genes activated by PP2A inhibition showed significant association with KRAS upregulation gene signature, indicating a novel transcriptional layer for PP2A-mediated antagonism of RAS signaling (Fig. 5C). Other significant cancer associated pathway signatures upregulated upon PP2A inhibition included epithelial mesenchymal transition (EMT), and mitotic spindle (Fig. 5C). Genes downregulated by PP2A inhibition were instead not enriched to any cellular process, supporting the conclusions that PP2A inhibition conveys its oncogenic effects primarily by gene activation. On the other hand, GO analysis of PP2A inhibited cells showed an enrichment of “positive regulation of intracellular signal transduction” indicating that in addition to its direct role in protein dephosphorylation, PP2A regulates signaling pathway activities transcriptionally. Other significant GO terms indicated the role for PP2A in regulation of cellular adhesion, migration and motility, all highly relevant for malignant cancers (Fig. S6). GSEA of genes downregulated by RAS inhibition, revealed an overlap with PP2A-regulated GSEA signatures, such as inflammatory response, and mitotic spindle (Fig. 5C, D). In addition, RAS was found to regulate genes related to IL6-JAK-STAT signaling and G2M checkpoint (Fig. 5D). On the other hand, the most significantly enriched GO biological process upon RAS silencing was “negative regulation of cell proliferation activity” (Fig. S6).

Notably, the convergence of RAS and PP2A activities on oncogenic transcription was also apparent via analysis of enrichment of transcription factor binding motifs on differentially regulated genes. TEAD (TEAD1, TEAD4 and YAP) and FOS (FOS, FOSL2) target genes were significantly enriched among those regulated by both PP2A and RAS targeting (Fig. 5F, in bold). This is consistent with recent results that PP2A complexes containing Striatins stimulate YAP1 activity leading to cellular transformation (Kurppa & Westermarck, 2020). On the other hand, YAP1 drives resistance to KRAS and EGFR inhibition (Kurppa, Liu et al., 2020, Shao, Xue et al., 2014). Further, KRAS and YAP1 synergistically activate FOS transcription factors, leading to EMT (Shao et al., 2014). In addition, AR target genes were also enriched among PP2A and RAS regulated targets (Fig. 5F, in bold). This is consistent with previous results indicating role for both RAS activity and PP2A inhibition in promoting malignant growth of AR-positive prostate cancers (Khanna, Rane et al., 2015, Weber & Gioeli, 2004).

Collectively, the RNA-seq data further supports the conclusions that PP2A inhibition drives oncogenic transcription, and that PP2A and RAS activities converge on transcriptional regulation of gene expression at various levels.

### DNA hypomethylation in PP2A-inhibited cells

Since both the PP2A-regulated phosphoproteome targets (Fig. 2A), and increased activity of the methylation-sensitive reporter assay (Fig. 4), indicated that PP2A could regulate DNA methylation, we analyzed global DNA methylation in PP2A inhibited cells by Reduced Representation Bisulfite Sequencing (RRBS). Consistent with the transcriptional de-repression (Fig. 4 and 5), the siRNA mediated inhibition of PP2A-A predominantly resulted in DNA demethylation (Fig. 6A). Of the total 211 differentially methylated regions, 143 regions showed a decrease in methylation marks (hypomethylated), while 68 showed an increase in methylation (hypermethylated) (Fig. 6B). The occupancy of the differentially regulated methylation marks was lowest at the exons (7%), while almost symmetrically distributed between introns (36%), intergenic areas (30%), and promoter regions (27%) (Fig. 6C). Inhibition of PP2A had an overall maximum impact on methylation of chromosome 11 (Fig. 6D). On the other hand, when the ratio between hypomethylation and hypermethylation was considered, the highest degree of hypomethylation was seen in X chromosome, which was exclusively hypomethylated, followed by the chromosome 13 in which six regions were hypomethylated and only one region was hypermethylated in response to PP2A inhibition (Fig. 6D).

**Figure 4.**
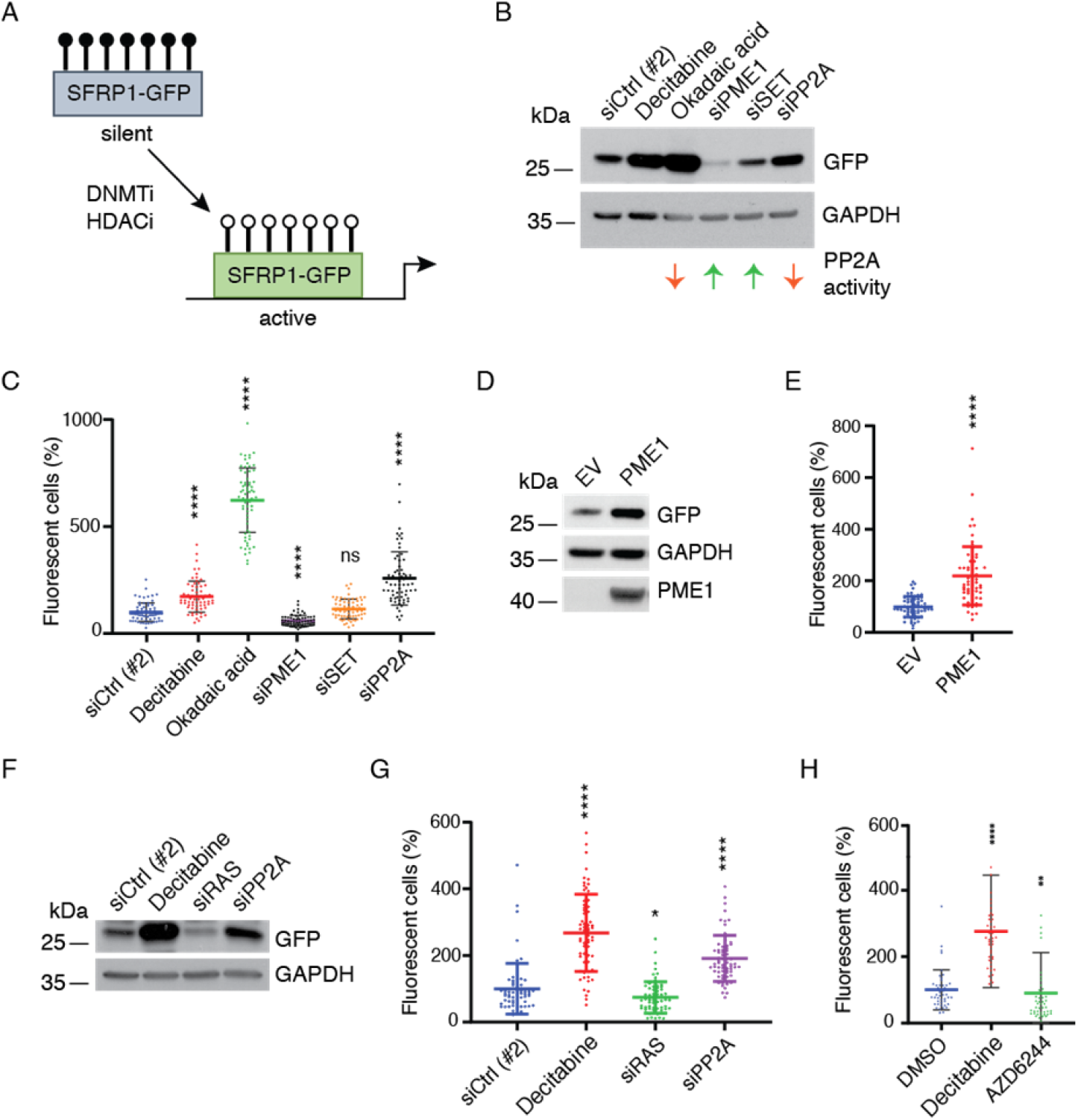
Opposing roles for PP2A and RAS in transcription from an epigenetically silenced SFRP1 promoter. **A**. Schematic representation of the SFRP1-GFP reporter system, which is activated by treatments that de-repress gene silencing by reversing DNA hypermethylation. The system has been previously shown responsive to DNMT1 and HDAC inhibitors (Cui et al., 2014). **B**. Western blot of reporter cells after treatment with Decitabine (DNMTi) as a control, and PP2A activation or inhibition **C**. Analysis of fluorescence in reporter cells after the treatments. **D**. Western blot of reporter cells after the overexpression of the PP2A inhibitor PME1. **E**. Analysis of fluorescence in reporter cells after the overexpression of PME1. **F**. Western blot of cells after treatment with Decitabine as control, and PP2A or RAS inhibition. **G**. Analysis of fluorescence in reporter cells in the same conditions. **H**. Analysis of fluorescence in reporter cells upon treatment with Decitabine and AZD6244 (MEKi).

**Figure 5.**
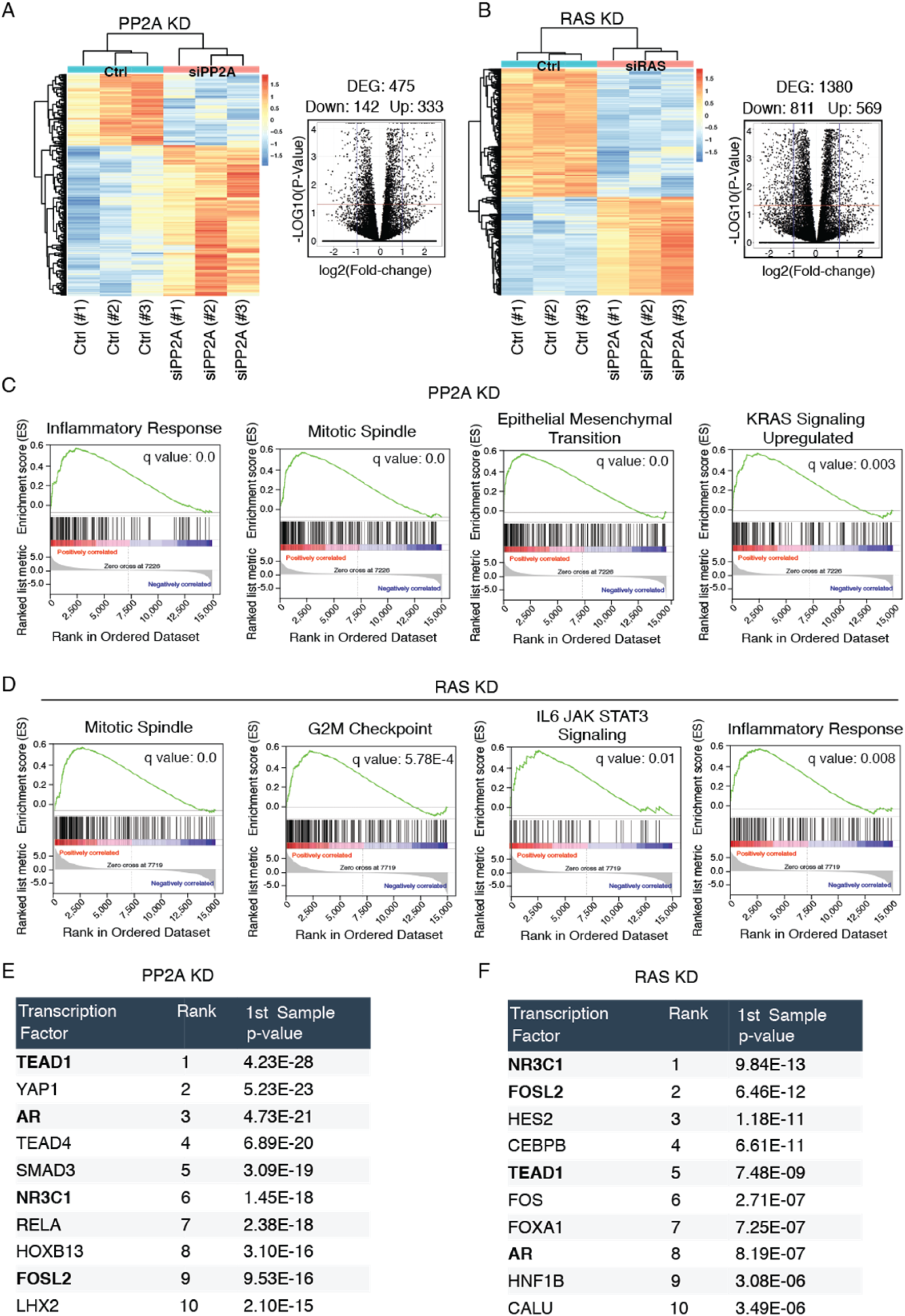
Converge of PP2A and RAS activities on oncogenic gene expression. **A**. Heat map of RNA-seq analysis of cells after PP2A inhibition and the corresponding volcano plot showing differentially regulated genes. **B**. Heatmap of RNA-seq analysis of cells after RAS inhibition and the corresponding volcano plot showing differentially regulated genes. **C**. Enrichment plots from gene set enrichment analysis (GSEA) of PP2A inhibited genes. **D**. GSEA enrichment plots of RAS inhibited genes. **E**. Top ten transcription factors regulated upon PP2A silencing. **F**. Top ten transcription factors regulated upon RAS silencing. The overlapping transcription factor elements enriched in both PP2A and RAS targets are bolded.

**Figure 6.**
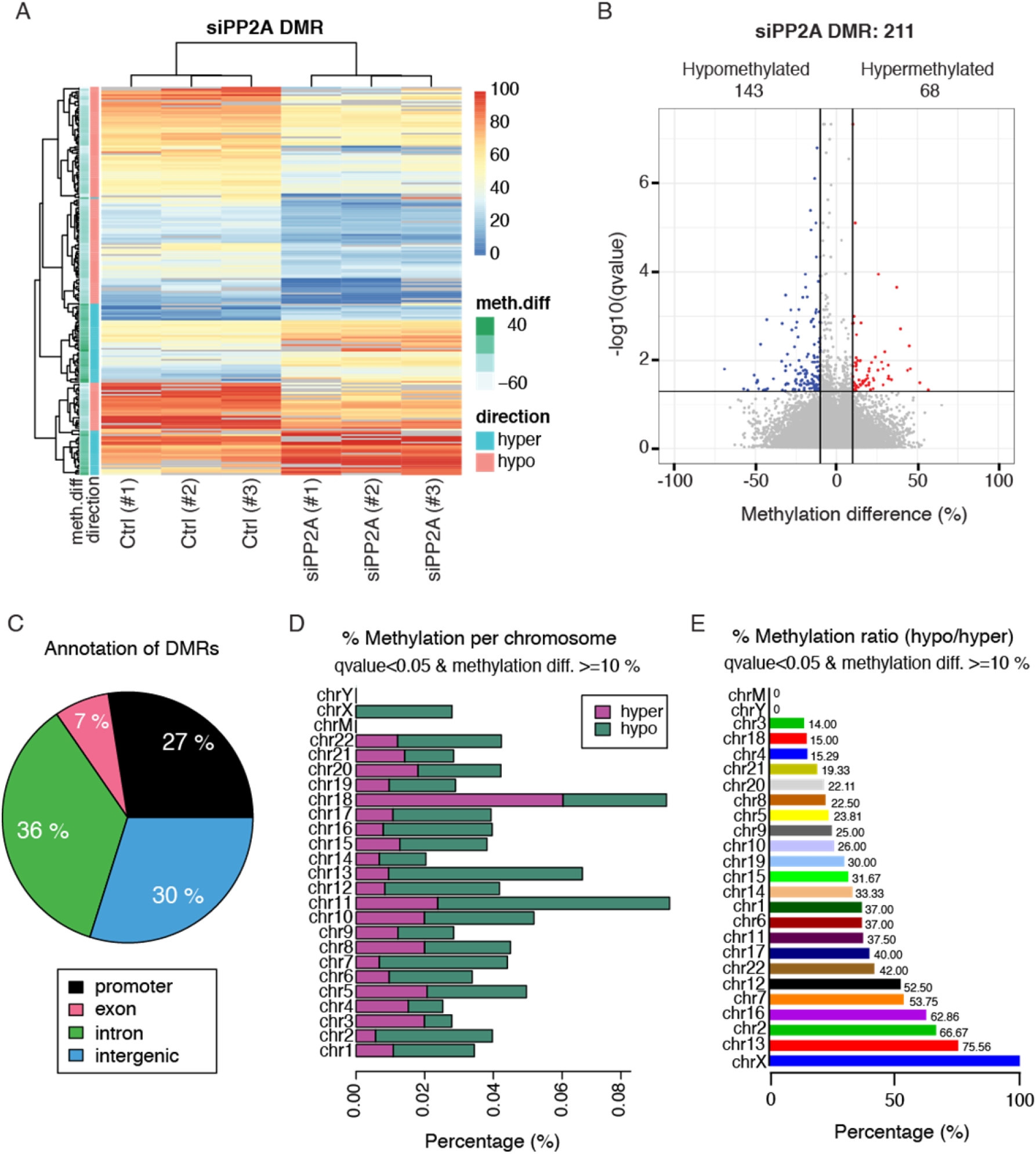
PP2A inhibition promotes DNA de-methylation. **A**. Heatmap showing the sample grouping according to CpG methylation levels (red = 100% methylated, blue = 0 % methylated9) upon PP2A inhibition. **B**. Volcano plot for differentially methylated regions, the blue dots indicate the significantly regulated hypomethylated regions while the red dots hypermethylated. **C**. A pie chart showing the percentage distribution of DMR among the genomic regions. **D**. Bar graph showing the distribution of DMR’s across different chromosomes. **E**. Bar graph depicting the ratio of hypo to hyper methylated regions among the chromosomes in ascending order.

To interrogate PP2A function in oncogenic transcription via its impact on DNA methylation, we performed a GO enrichment analysis based on differentially methylated regions (DMRs), and by using the Enrichr analysis tool (Kuleshov, Jones et al., 2016) (Fig. S7). Importantly, overlapping with the GSEA analysis of PP2A-regulated gene expression (Fig. 5C), both epithelial to mesenchymal transition, and several gene sets related to membrane associated GTPase activity (corresponding to KRAS activity in GSEA), were enriched in PP2A inhibited cells (Fig. S7).

These results provide novel evidence for PP2A-mediated regulation of DNA methylation, and indicate that DNA hypomethylation contributes to PP2A inhibition-induced oncogenic transcription.

### PP2A regulates chromatin accessibility

The PP2A phosphoproteome targets (Fig. 2A) indicated that PP2A may regulate gene expression also by affecting chromatin accessibility. This prompted us to perform Assay for Transposase-Accessible Chromatin using sequencing (ATAC-seq) analysis from cells transfected with control and PP2A A-subunit siRNAs. Supporting our hypothesis, PP2A inhibition induced marked changes in the pattern of open chromatin regions (Fig. 7A, B). The number of open peaks upon PP2A silencing were 9,437, and the general distribution resembled that of ATAC-seq profiles in general (Yan, Powell et al., 2020) (Fig. 7A). The most enriched binding sites for transcriptional regulators associated with genes with open and closed promoter regions are listed in panels 7D and 7E, respectively. Very interestingly, PP2A and RAS phosphotarget DOT1L (Fig. 2B) was among the regulators associated with open promoter regions (Fig. 7D), providing a direct link between the phosphoproteome and ATAC-seq data. Additionally, the differently accessible peaks in PP2A-inhibited cells were found in proximity to genes involved in chromosome and chromatin binding, indicating for another putative layer of regulation of how PP2A inhibition may impact oncogenic transcription.

**Figure 7.**
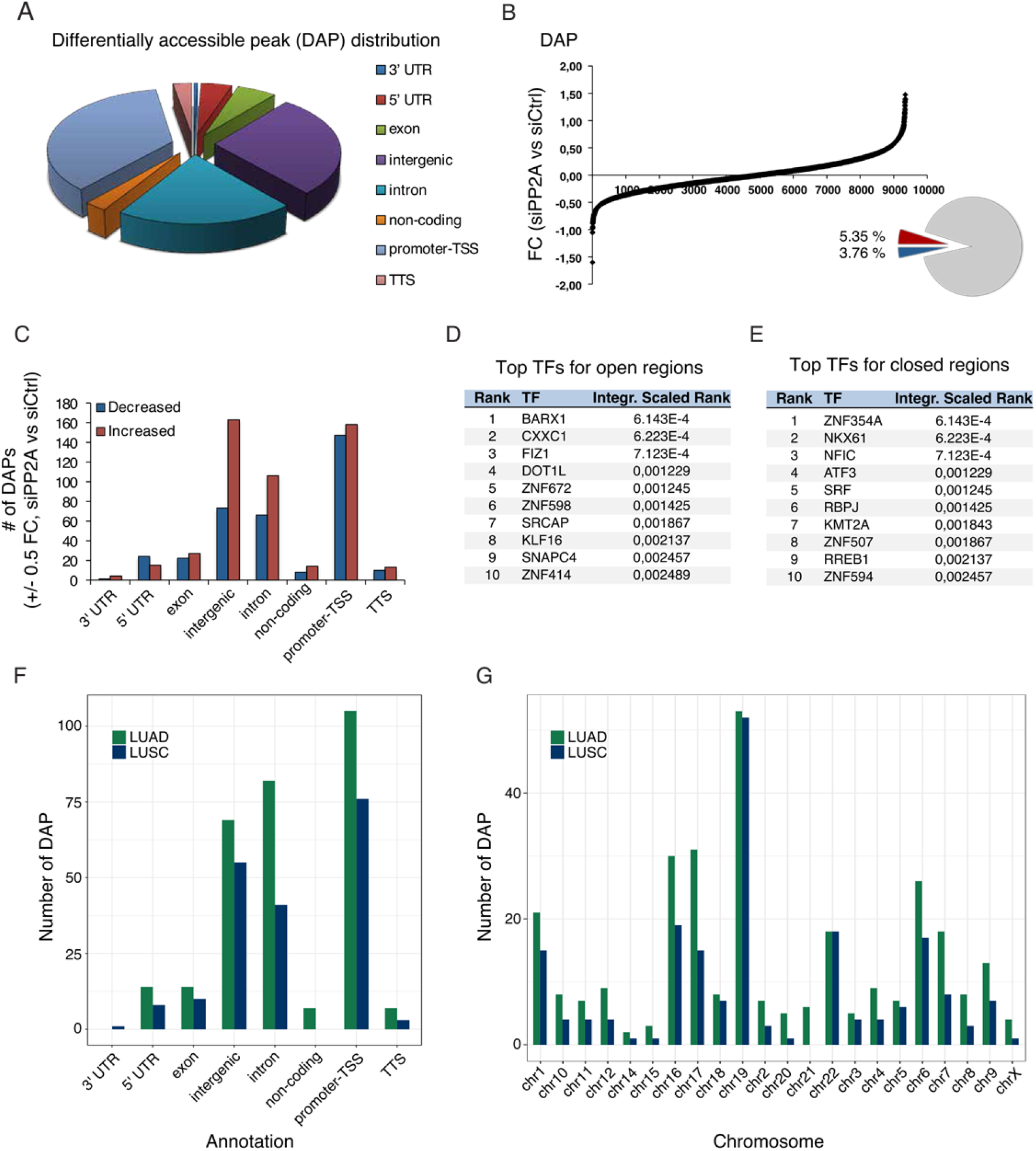
The chromatin re-modelling upon PP2A inhibition and its overlap to clinical patient sample chromatin landscapes. **A**. The pie chart shows the distribution of differentially accessible peaks of the experiment. The open regions/peaks per genomic area resembles that of ATAC-Seq. profiles in general. **B**. The graph illustrates the total number of open/closed areas that has changed upon PP2A transient knockdown as compared to siCtrl. Of the identified peaks 5.35 % are opened and 3.76 % closed (FC +/-0.5) in siPP2A vs siCtrl. **C**. The chart shows the distribution of these areas per genomic area, indicating that most changes occur in the promoter-TSS area, while there is more open than closed areas upon siPP2A in intergenic areas. **D**. Genes in proximity to promoter-TSS are regulated by these top ten transcription factors. **E**. The graph shows the transcription factors binding to the genes in proximity to promoter-TSS within the closed regions upon PP2A inhibition. The graph shows the top ten transcription factors. **F**. The graph shows the differentially accessible peaks upon siPP2A that are overlapping with clinical patient material per genomic area **G**. Shows the differential accessible overlapping peaks of siPP2A that are overlapping with clinical patient material as distributed per chromosome.

Next, we were interested to know whether the observed PP2A-dependent modulation of chromatin accessibility could have a clinical relevance. Therefore, we compared the ATAC-seq profiles from the PP2A inhibited cells to NSCLC patient ATAC-seq profiles, including lung adenocarcinoma (LUAD) and lung squamous cell carcinoma (LUSC) (Wang, Tu et al., 2019). Interestingly, a number of open chromatin peaks identified in the PP2A inhibited cells were found to overlap with open chromatin peaks in NSCLC patient samples with highest degree of overlap in chromosome 19 (Fig. 7F, G).

Collectively these results provide novel evidence that PP2A inhibition impacts chromatin accessibility.

## DISCUSSION

Epigenetic gene regulation has an established role in cancer initiation and progression (Baylin & Jones, 2016, Cheng et al., 2019, Laugesen & Helin, 2014, Quagliano et al., 2020). Several loss-of-function studies across different species have also demonstrated an integral role for epigenetic gene regulation in signal transduction, development, and malignant progression downstream of RAS proteins (Vaz et al., 2017). However, it has been surprisingly poorly known how RAS impacts phosphorylation of epigenetic proteins, and whether RAS activity towards epigenetic gene regulation is modulated by PP2A-mediated protein dephosphorylation. Here we provide the first bird’s-eye view to the global impact of RAS and PP2A activities on the phosphorylation regulation of epigenetic complexes and their co-operative impact on oncogenic gene regulation. The results indicate that epigenetic protein complexes involved in oncogenic gene expression constitute a significant point of convergence for RAS hyperactivity and PP2A inhibition in cancer. The results also provide a very rich resource for future interrogation of the impact of the identified phosphosites both in physiological gene regulation, as well as in human cancer development and progression.

About 20 years ago, inhibition of serine/threonine phosphatase activity of PP2A was established by several studies as a prerequisite for RAS-mediated malignant transformation of human, but not of mouse, cells (Hahn et al., 2002, Rangarajan et al., 2004, Yu et al., 2001). Mechanistically the requirement of PP2A inhibition for RAS-mediated human cell transformation was attributed to the role of PP2A as an inhibitor of the activities of several downstream mediators of RAS activity such as MEK/ERK and AKT kinases, or transcription factor MYC (Sablina et al., 2010, Yeh, Cunningham et al., 2004). In this prevailing model, PP2A inhibition is seen merely as a mechanism to boost signal transduction initiated by RAS. However, it has not been previously systematically addressed whether their activities would converge on particular cellular mechanisms or processes. This is a critical unanswered question, as understanding why RAS activity and PP2A inhibition are mutually required for human cell transformation could lead to fundamental novel understanding of the basis of human cancer development and facilitate novel therapeutic approaches that target the roots of human malignancies. In this context, PP2A inhibition has been demonstrated to drive resistance of KRAS mutant cells towards a wide array of kinase inhibitors (Kauko et al., 2018). Further, we and others have shown that PP2A inhibition drives cancer cell resistance to epigenetic therapies (Fig. S5D-F) (Kauko et al., 2020, Shu, Lin et al., 2016). Thereby the presented results may provide important clues for understanding the role of PP2A-mediated phosphoregulation on responses of RAS-driven cancers to epigenetic therapies that has thus far been clinically disappointing. On the other hand, the results encourage testing of the emerging PP2A reactivating therapies (Vainonen, Momeny et al., 2021) for their impact on KRAS-mutant cancers in combination with epigenetic therapies.

Our data provides strong evidence that PP2A inhibition drives oncogenic transcription and globally impacts both DNA methylation and chromatin accessibility. While the role of PP2A inhibition in oncogenic transcription is fully consistent with its role in the CDK9-mediated (RNAPII-driven) transcriptional elongation (Huang et al., 2020, Vervoort et al., 2021), the role of PP2A in DNA methylation and chromatin accessibility has been largely uncharacterized. While DNA methylation of the promoters is the most studied epigenetic mechanism regulating gene expression, recent reports indicate that cancer cells harbor hypomethylated regions at the intergenic regions (Lee & Wiemels, 2016). Thereby the observed hypomethylation at intergenic regions by PP2A inhibition indicate novel possible regulatory mechanisms for cancer progression. On the other hand, our results demonstrating global chromatin opening by PP2A inhibition are consistent with recent results from mouse T cells in which deletion of regulatory B-subunit of PP2A (PPP2R2D) resulted in chromatin opening (Pan, Sharabi et al., 2020).

Our results reveal dozens of RAS- and PP2A-regulated phosphorylation sites in epigenetic proteins previously implicated in transcription and cancer (Fig. 2, and S2). Structurally we provide evidence that at least some of these phosphosites are located on functionally important regions of the epigenetic proteins involved in oncogenic transcription (Fig. 2C). Mechanistically we validate the impact of RAS and PP2A on selected NuRD complex members. While the mechanism by which RAS and PP2A regulate CHD3 protein stability remains speculative, loss of CHD3 expression upon RAS inhibition (Fig. 3B) most likely contributes to the observed chromatin remodeling phenotype in RAS/PP2A-modulated cells (Fig. 7). On the other hand, previous studies have shown that HDAC2 phosphorylation is required for its interactions with epigenetic multiprotein complexes such as Sin3, NuRD or CoREST (Delcuve, Khan et al., 2012). In our data both RAS and PP2A regulate C-terminal HDAC1 and HDAC2 phosphorylation on largely overlapping sites (Fig. 2). Functionally we validate that RAS and PP2A modulation regulate chromatin binding of HDAC1/2 and that this correlates with transcriptional activity of highly HDAC responsive SFRP1 promoter system. Naturally both phenotypes can also be attributed to RAS/PP2A-mediated phosphorylation regulation of other NuRD complex components such as MTA2. Additionally, we cannot exclude that RAS and PP2A regulates gene expression and chromatin accessibility by directly regulating histones or transcription factors (Gil & Vagnarelli, 2019). Interestingly, in addition to RAS/PP2A-mediated phosphoregulation of epigenetic proteins reported here, PP2A has been shown to dephosphorylate BRD4, HDAC 4/5/7, PRMT1/5 and TET2 that can contribute to chromatin structure regulation (Tinsley & Allen-Petersen, 2022) Therefore, these data collectively indicate that precise control of gene expression relies on the finely tuned balance between kinase and phosphatase activities.

Collectively these data reveal a previously hidden layer of phosphoregulation of epigenetic gene regulation. Based on the results it is very likely that convergence of the RAS and PP2A activities on the discovered epigenetic phosphoregulation contribute also to the synergism of RAS activation and PP2A inhibition in human oncogenesis. We further postulate that the discovered RAS/PP2A-mediated phosphorylation switches are most probably not only relevant in cancer, but also in development and other diseases.

## ACKNOWLEGDEMENTS

We thank The Proteomics and The Cell Imaging Core at Turku Centre for Biotechnology supported by the University of Turku and Biocenter Finland, for technical assistance. Taina Kalevo-Mattila is acknowledged for excellent technical help and the entire Turku Bioscience Centre technical staff for their contributions. This work was supported, in part, by grants from Academy of Finland (JW, 294850), and Jane ja Aatos Erkko Foundation (JW). MS has been supported by the Turku Doctoral Programme of Molecular Medicine (TuDMM), Ida Montin Foundation and the Paulo Foundation. We acknowledge the following colleagues for generously providing valuable research tools as listed in the materials and methods: Prof. Krister Wennerberg, Prof. Gautham Narla, Prof. Jerry W Shay, Prof. Stephen B. Baylin and Dr. Michael Ohlmeyer. The Structural Genomics Consortium is a registered charity (no: 1097737) that receives funds from Bayer AG, Boehringer Ingelheim, Bristol Myers Squibb, Genentech, Genome Canada through Ontario Genomics Institute [OGI-196], EU/EFPIA/OICR/McGill/KTH/Diamond Innovative Medicines Initiative 2 Joint Undertaking [EUbOPEN grant 875510], Janssen, Merck KGaA (aka EMD in Canada and US), Pfizer and Takeda.

## MATERIALS AND METHODS

### Phosphoproteome data filtering

The details of the phosphoproteomics pipeline and data analyses related to analyses of RAS and PP2A-regulated phosphosites are described in previous publications (Kauko et al., 2020, Kauko et al., 2015). The raw data can be accessed via the PRIDE partner repository with the dataset identifier PXD001374 (for RAS regulated phosphosites) and PXD016102 (for PP2A-regulated phosphosites). For identification of overlapping phosphosites regulated by RAS, or any of the PP2A conditions, the following filtering criteria was used when assessing the phosphoproteome data normalized as described in (Kauko et al., 2020, Kauko et al., 2015). For inhibition of phosphorylation by RAS targeting: fold change -0.5 log2 and FDR<0.1%. For PP2A-regulated proteins fold change 0.5 log2 (for increased phosphorylation by PPP2R1A targeting) and -0.5 log2 (for dephosphorylation by PME-1, CIP2A and SET targeting) and FDR<0.05%. The lower FDR criteria used for RAS data is due to notion that in general these earlier experiments (Kauko et al., 2015) had more variation between the replicate samples due to less sensitive mass spectrometry and inexperience in sample handling.

### Cell culture

KRAS mutant lung cancer cell lines A549, a kind gift of Prof. Krister Wennerberg, and H358, a kind gift of Prof. Gautham Narla, H460 as well as HeLa cells were used. Furthermore, the normal bronchial epithelial cell line HBEC3-KT (HBEC), immortalized with CDK4 and hTERT, a kind gift of Prof. Jerry W Shay (Sato et al., 2013), was used. HCT116 SFRP1 promoter-GFP cells (Cui et al., 2014) were a kind gift of Prof. Stephen B. Baylin. A549, H358, H460 and HBEC cells were authenticated (STR profiling) by the European Collection of Authenticated Cell Cultures (ECACC) in December 2018. The cells were cultured in medium conditions recommended by the providers for less than 4 months before use in these experiments. All cells were regularly tested negative for mycoplasma.

### siRNA transfections and treatments

The siRNA transfections were done in 6 well-to 96-well plates, utilizing RNAiMAX transfection reagent according to manufacturers instructions, and siRNAs from Qiagen and Eurofins.

siRNA sequences and/or provider cataloge #:

siCtrl (#1): AllStars Neg. Control siRNA, Cat. No. / ID: 1027281 (Qiagen), siCtrl (#2): CGUACGCGGAAUACUUCGA (Eurofins), siPP2A-A: UUUUCCACUAGCUUCUUC A (Eurofins), siHRAS: GAACCCUCCUGAUGAGAGU (Eurofins), siKRAS: AGAGUGCCUUGACGAUACA (Eurofins), siNRAS: GAAAUACGCCAGUACCGAA, siPME (#1): GGAAGUGAGUCUAUAAGCA, siPME1 (#2): UCAUAGAGGAAGAAG AAG A, siSET (#1): UGCAGACACUUGUGGAUGG (Eurofins), siSET (#2): AAUGCA GUGCCUCUUCAUC (Eurofins). All siRNAs (CHD3, DNMT1, DOT1L, KDM1A, MLLT3, RNF168 and SMARCA4) for the cell viability assay were ordered from Qiagen. AllStars Hs Cell Death siRNA, Cat. No. / ID: 1027299 (Qiagen), was used as a positive control (siCtrl +).

DNMT1 inhibitors (AZA, Decitabine), BET inhibitors (iBET-151, JQ1, Mivebresib), HDAC inhibitors (Panobinostat, TSA), KDM1A inhibitors (SP2509) and Ocadaic acid were used, and purchased from SelleckChem. PP2A reactivating compound DBK1154 was a kind gift of Dr. Michael Ohlmeyer.

### Nuclear fractionation

Cell fractionation was done using the Subcellular Protein Fractionation Kit for Cultured Cells from Thermo Scientific (#78840). Briefly, cells were harvested 48 h post transfection using trypsin. One million cells were suspended in 100 µl of cytoplasmic extraction buffer and incubated for 10 min at 4°C with gentle mixing. Cells were then centrifuged at 400g for 5 min and the supernatant was collected as the cytoplasmic fraction in a new tube. Membrane extraction buffer (100 µl) was added to the pellets followed by vigorous vortexing for 5 sec and incubation at 4°C for 10 min. The lysate was centrifuged at 5000g for 5 min, and the supernatant containing the membrane extracts was collected in a fresh tube. The pellet was suspended in the nuclear isolation buffer (50 µl) and vortexed for 15 seconds and further incubated for 30 min at 4°C with gentle rotation. Cells were centrifuged at 5000g for 5 min and the supernatant containing the nuclear fractions was collected. The pellets were suspended in nuclear isolation buffer (50 µl) containing MNAse (150 U) and 5 mM Calcium chloride and vortexed for 15 sec. The tubes were incubated at RT for 15 min to separate the chromatin bound proteins. After vortexing again for 15 sec tubes were centrifuged at 16000g for 5 min and the supernatant containing the chromatin bound proteins was collected in fresh tubes. Protein concentration of the fractions was determined using the BCA assay and Western blotting was used to detect cellular localization of the desired proteins.

### Pull-down assays

To confirm the interaction between PP2A-B56α and HDAC1 pull down assay was performed as described before (PMID: 34145035). Briefly, the cells were transfected with the respective plasmids using the jetPRIME® transfection reagent. 48 h later cells were harvested on ice by scraping and lysed in a buffer containing 100 mM NaCl, 1 mM MgCl2, 10 % glycerol, 0.2 % protease inhibitor tablet (Roche) and 25 units/ml Benzonase (Millipore). Cells were rotated at 4°C on a roller and fifteen min later the final concentration of NaCl and EDTA was increased to 200mM and 2mM, respectively. After further rotation of 10 min cells were centrifuged at 16,000 rpm for 20 min. 10% of the lysate was stored as input and the remaining was incubated with the 20µl of prewashed GFP trap magnetic beads (ChromoTek GFP-Trap®) at 4°C on a roller for two hours. Post incubation the beads were washed three times using the lysis buffer and eluted by adding 20 µl of 2X SDS loading buffer and boiling at 95°C for 10 min. Inputs and the CO-IP samples were further loaded on a 4-20% gradient gel to access the interactions.

### Western blots

Cells were lysed in RIPA buffer (50 mM Tris-HCl pH 7.5, 0.5 % DOC, 0.1 % SDS, 1% NP-40, and 150 mM NaCl) with protease and phosphatase inhibitors (#4693159001 and #4906837001, Roche) followed by sonication at highest setting with a pulse of ± 30 sec. After centrifugation at 16,000g for 30 min lysates were collected in a fresh tube and protein concentration was determined using BCA assay (Pierce). 6X loading buffer was added to lysates and they were boiled at 95°C for 10 min. Equal amounts of lysates were load in 4-20% precast gradient gels (Bio-Rad) and separated at 80-100 Volts. Proteins were blotted using PVDF membrane (Bio-Rad) and blocked for 1 h at RT. Membranes were incubated overnight with primary antibody followed by washing. For detection, HRP-labelled secondary antibodies (DAKO) followed by incubation with Pierce™ECL Western Blotting Substrate (Thermo Fisher Scientific) was used, or LI-COR Biosciences secondary antibodies (IRDye 680 or IRDye 800) was used followed by detection by Odyssey® Imaging Systems or Bio-Rad Laboratories ChemiDoc Imaging Systems.

### Antibodies

The following antibodies, at the indicated dilutions, were used:

FLAG: Sigma-Aldrich F3165 (1:1000), GAPDH: Hytest 5G4-6C5 (1:5000), GFP:

Santa Cruz Biotechnology sc-9996 (1:500); HDAC1 and 2: Sigma 06-720-25UG and Santa Cruz Biotechnology sc-9959, respectively (1:1000), H3: Santa Cruz Biotechnology sc-374669 (C-2) (1:1000), PME1: Santa Cruz Biotechnology sc-20086 (H-226) (1:1000), SET1: I2PP2A (F-9) Santa Cruz Biotechnology sc-133138 (1:1000).

### Drug sensitivity assays

To determine the synergy between PP2A activation and HDAC inhibition drug synergy screen was done. H460 cells were seeded in 96 well plate (3000 cells/well) and next day treated with respective drugs for 48 h. Cell viability was measured using the CellTiter-Glo® cell viability end-point assay (Promega), and the synergy was determined using the synergy finder tool (https://synergyfinder.fimm.fi/synergy/20220404143330175356/). HCT116 reporter cells were treated with the respective drugs at their IC50 concentrations, and 48 h later imaged for fluorescence using the IncuCyte ZOOM and/or S3 live cell imaging, and then harvested using RIPA buffer. Fluorescence signal was analyzed using the ImageJ tool while the GFP signal was determined using Western blotting.

### Anchorage-independent colony formation assay

For the anchorage-independent colony formation assay, which typically correlates with *in vivo* tumorigenicity, 2×10^4^ cells were resuspended in 1.5ml growth medium containing 0.4% agarose (4% Agarose Gel, Termo Fisher Scientific Gibco; top layer) and plated on 1ml bottom layer containing growth medium and 1.2% agarose in a 12-well plate. After 14 days of growth, colonies were stained over night with 1mg/ml Nitro blue tetrazolium chloride (NBT; Molecular Probes) in PBS. Colonies were imaged using a Zeiss SteREO Lumar V12 stereomicroscope. Analysis was done using the ImageJ software. First, the background was subtracted using the rolling ball function with a radius of 50μm, then auto-thresholding was applied to separate the colonies. Area percentage was calculated using the ImageJ built-in function ‘Analyze Particles’ with exclusion of particles smaller than 500μm^2^ that are not considered colonies.

### RNA-sequencing

HeLa cells were transfected with the siRNAs using Lipofectamine RNAiMAX. After 72 h RNA, DNA & Protein were isolated using the AllPrep DNA/RNA/Protein Mini Kit (50) Qiagen Cat. No. / ID: 80004). RNA sequencing was done at Finnish Functional Genomics Centre (Turku, Finland). First the quality of the total RNA samples was ensured with Advanced Analytical Fragment Analyzer. Sample concentration was measured with Qubit® Fluorometric Quantitation, Life Technologies. Library preparation was done according to Illumina TruSeq® Stranded mRNA Sample Preparation Guide (part # 15031047). The samples were sequenced with Illumina HiSeq 3000 instrument using single-end sequencing with 1 × 50 bp read length. RNA-seq data quality was assessed with FastQC v0.11.7 (“FastQC,” 2015, retrieved from https://qubeshub.org/resources/fastqc)•. Reads were trimmed with TrimGalore! v0.6.4._dev (https://zenodo.org/record/5127899)• with the following parameters: -- quality 20 --gzip -fastqc. Resulting trimmed single-end reads were aligned with STAR v2.5.3a with the following parameters: --quantMode TranscriptomeSAM -- outSAMtype BAM SortedByCoordinate --chimOutType WithinBAM –twopassMode Basic --readFilesCommand zcat --genomeLoad NoSharedMemory -- outReadsUnmapped FastX ---bamRemoveDuplicatesType UniqueIdentical -- seedSearchStartLmax 25 --outFilterMismatchNoverReadLmax 25 -- outFilterMismatchNoverReadLmax 0.04 --winAnchorMultimapNmax 100. Ensembl Homo Sapiens GRCh38 v95 sequence and annotations were used as reference for the alignment. Gene counts produced with STAR were then assembled into a read count matrix within R v4.1.0 (R Core Team, 2019, retrieved from https://www.r-project.org/)• and DESeq2 v1.34.0 (PMID: 25516281)• was used to detect differentially expressed genes. For each comparison, pre-filtering was applied to the corresponding data matrix: genes with zero read counts across all samples were removed, the lower quartile value of the resulting distribution was computed and genes with overall read count lower than this value removed. DEseq was run with default parameters. GO (PMID: 10802651)• terms enrichment was computed with goseq v1.46.0 (PMID: 20132535)• using non-electronic GO associations (IEA associations were removed).

### RRBS

The DNA isolated from the same samples was used for RRBS sequencing. RRBS was done at Finnish Functional Genomics Centre (Turku, Finland). Initially the quality of the genomic DNA samples was ensured with Advanced Analytical Fragment Analyzer and concentrations were measured with Qubit® Fluorometric Quantitation, Life Technologies. Library preparation was carried out according to protocol adapted from Boyle et al. (Boyle, Clement et al., 2012). Bisulfite conversion and sample purification were done according to Invitrogen MethylCode Bisulfite Conversion Kit. The samples were sequenced with Illumina HiSeq 3000 instrument using single-read sequencing with 1 × 50 bp read length.

RRBS data quality was assessed with FastQC v0.11.8 (“FastQC,” 2015, retrieved from https://qubeshub.org/resources/fastqc)•. Reads were trimmed with TrimGalore! v0.6.4_dev (https://zenodo.org/record/5127899)• (running on top of Cutadapt v2.7 (https://doi.org/10.14806/EJ.17.1.200)•) with the following parameters: --quality 22 -- phred33 --gzip --rrbs --fastqc --paired --cores 4. Resulting trimmed paired-end reads were aligned with Bismark v0.22.3 (PMID: 21493656)• using the Bowtie2 aligner v2.3.5.1 (PMID: 30020410)• and the following additional parameters: --unmapped -- ambiguous --ambig_bam --nucleotide_coverage --fastq. Methylation calls were extracted with the bismark_methylation_extractor and the following parameters: -- paired-end --comprehensive --gzip --bedGraph --remove_spaces --buffer_size 80% - -cytosine_report --ignore_r2 2. Data were aligned to the Ensembl Homo Sapiens GRCh38 v95 genome and corresponding annotations were used. Incomplete conversions were filtered out with the filter_non_conversion tools from Bismark. Finally, differential methylation analysis was run using Bioconductor (Huber, Carey et al., 2015) • methylKit v1.12.0 (Akalin, Kormaksson et al., 2012) • packages running on R v3.6.1 (R Core Team, 2019 retrieved from https://www.r-project.org/)•. For the methylKit analysis two scenarios were considered: CpG context and tiled window context. For single base differential methylation calls the following parameters were set: methylation difference 25%, adjusted q-value 0.01. On top of these, for the tiled analysis, window size and step size were set to 500 bp to generate tiling, non-overlapping windows. The whole analysis was bundled in a Snakemake pipeline and software specifications encapsulated in a Singularity container. The analysis was run on Snakemake v5.6.0 (Molder, Jablonski et al., 2021). Both the workflow and container recipies are available at https://github.com/ftabaro/MethylSnake.

### ATAC-sequencing

To profile the open chromatin regions, ATAC-seq was conducted, according to the protocol by Buenrostro and co-workers (Buenrostro, Wu et al., 2015). HeLa cells were transfected with siCtrl or siPP2A-A, and 72 h upon transfection 50,000 cells were collected for analysis. At the end the library sizes were determined by fragment analysis, and 2×75 paired-end sequencing was performed on NextSeq500 (Illumina) to yield an average of 50 M reads/sample. Sequencing libraries quality was assessed with FastQC v0.11.7. Reads were aligned on the human genome GRCh38 with Bowtie2 v2.3.4.1, converted to bam with samtools v1.8, sorted with Picard SortSam v2.27.1. Bam files were indexed with Picard BuildBamIndex v2.27.1 and duplicated reads were marked with Picard MarkDuplicates v2.27.1. Peaks were called with MACS2 v2.1.0 with the following parameters: --gsize hs –qvalue 0.05 – format bam –nomodel –bdg –call-summits. Peak summits were annotated with Homer v4.9 by running the annotatePeak.pl routine against hg38 annotation. Standardized peaks were computed starting from peaks summit using the bedtools slop command v2.27.1 with parameters: -g $GENOME -l 250 -r 249. Read counts of standardized peaks was computed from bam files using the bedtools coverage command v2.27.1. Standardized peaks read count was normalized using median of ratios normalization, and then converted to RPM. Finally, log2 fold change was computed from RPM values comparing signal intensities of corresponding genomic locations between siPP2A and siCtrl samples. Peaks with absolute fold change greater than 0.5 were considered differentially accessible.

### Statistical analyses

Statistical analyses were completed using GraphPad Prism, GraphPad Software (www.graphpad.com). The data were analyzed with the Mann–Whitney U test for significance (RNAi screens) or the Student standard t test.

